# Insight into the genetic architecture of back pain and its risk factors from a study of 509,000 individuals

**DOI:** 10.1101/384255

**Authors:** Maxim B Freidin, Yakov A Tsepilov, Melody Palmer, Lennart C Karssen, CHARGE Musculoskeletal Working Group, Pradeep Suri, Yurii S Aulchenko, Frances MK Williams

## Abstract

Back pain (BP) is a common condition of major social importance and poorly understood pathogenesis. Combining data from the UK Biobank and CHARGE consortium cohorts allowed us to perform a very large GWAS (total N = 509,070) and examine the genetic correlation and pleiotropy between BP and its clinical and psychosocial risk factors. We identified and replicated three BP associated loci, including one novel region implicating *SPOCK2*/*CHST3* genes. We provide evidence for pleiotropic effects of genetic factors underlying BP, height, and intervertebral disc problems. We also identified independent genetic correlations between BP and depression symptoms, neuroticism, sleep disturbance, overweight, and smoking. A significant enrichment for genes involved in central nervous system and skeletal tissue development was observed. The study of pleiotropy and genetic correlations, supported by the pathway analysis, suggests at least two strong molecular axes of BP genesis, one related to structural/anatomic factors such as intervertebral disk problems and anthropometrics; and another related to the psychological component of pain perception and pain processing. These findings corroborate with the current biopsychosocial model as a paradigm for BP. Overall, the results demonstrate BP to have an extremely complex genetic architecture that overlaps with the genetic predisposition to its biopsychosocial risk factors. The work sheds light on pathways of relevance in the prevention and management of LBP.

## MAIN

Back pain (BP) is a common debilitating condition with a lifetime prevalence of 40% and a very important socioeconomic impact^1,2^. According to the Global Burden of Disease 2016 study, it leads the list of disabling conditions in many parts of the world^3^. Known clinical risk factors for BP include age, female gender and raised body mass index^4^. The greatest risk for episodes of severe BP in population based studies is thought to be attributable to intervertebral lumbar disc degeneration (LDD) ^5^, though its predictive and diagnostic impact remains debated^6^. In the majority of episodes of BP the symptoms are transient; however, about 10% of those experiencing acute BP develop a chronic condition^2^ which places a great socioeconomic burden on society^7–9^.

There is a clear genetic predisposition to BP with estimates of heritability in the range of 30%-68%^10–13^. Similar or higher heritability estimates for LDD have been obtained^14,15^. Importantly, not only is there a phenotypic association between LDD and LBP but a genetic correlation between the two has been reported in twin studies (11%-13%)^10,16^, suggestive of shared genetic background. Twin studies have demonstrated that BP also shares an underlying genetic predisposition with several of its risk factors including depression and anxiety^17^, educational attainment^18^, obesity^19^ as well as with other pain conditions such as chronic widespread musculoskeletal pain^20^.

We recently performed a genome-wide association study (GWAS) for chronic BP (BP lasting longer than 3 months) from the interim release of the UK Biobank^21^ and from the Cohorts for Heart and Aging Research in Genomic Epidemiology (CHARGE) Musculoskeletal Working Group^22^ (total N = 158,000 individuals). Despite a large sample size and relatively well-defined phenotype, the study identified and replicated only three loci associated with chronic BP. This suggests that the genetic architecture of BP is extremely complex and far larger samples are required to make progress in the field.

In the present study we sought to expand the BP GWAS and explore the genetic associations with many of the biopsychosocial risk factors for BP. In brief, we examined 350,000 individuals of European ancestry from the UK Biobank in the discovery phase (91,100 cases and 258,900 controls) followed by a replication phase combining the UK Biobank participants of European, African and Asian ancestry not included in the discovery set, and data from the CHARGE cohorts (total N = 157,752). Post-GWAS analyses included the analysis of pleiotropy, genetic correlations and pathway analyses (Figure 1).

**Figure 1.**
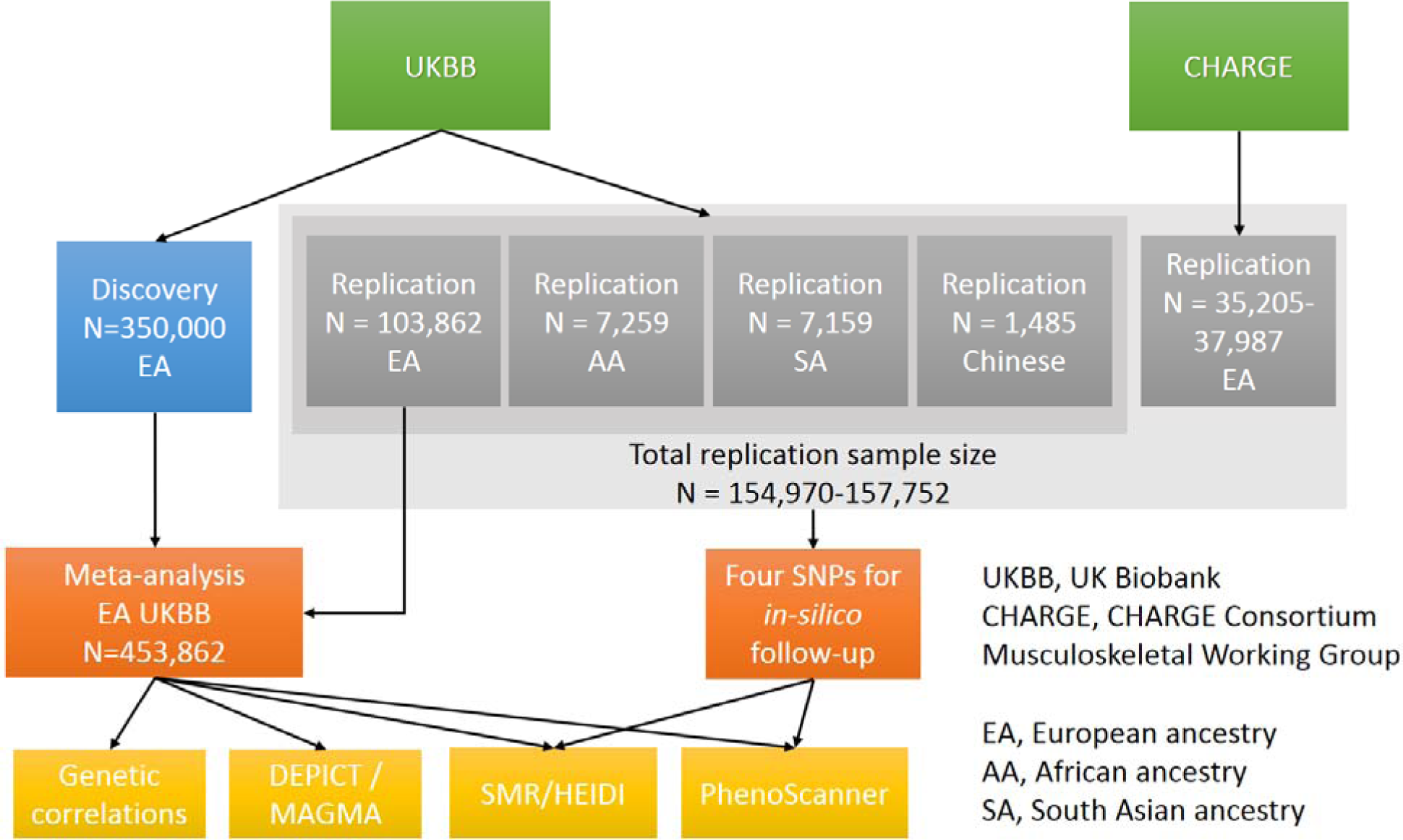
Overview of the study. GWAS for back pain used a combination of UK Biobank and Cohorts for Heart and Aging Research in Genomic Epidemiology (CHARGE) consortium cohorts. Discovery was performed using 350,000 individuals of European ancestry from the UK Biobank. Replication cohorts included individuals of European (EA), African (AA) and South Asian (SA) ancestry and Chinese individuals from the UK Biobank and CHARGE cohorts (N = 154,970-157,752). Meta-analysis was carried out using the discovery cohort and other individuals of European ancestry from the UK Biobank (N = 453,862) and the results used to estimate genetic correlations with risk factors, establish causal or pleiotropic relationships using summary-data based Mendelian randomization (SMR) followed by heterogeneity in dependent instruments (HEIDI) analysis, and to perform DEPICT and MAGMA analyses to reveal functional relevance.

## RESULTS

### Novel genomic loci associated with back pain

The discovery sample of white British individuals (as defined by genetic principal components; N = 350,000) comprised 91,100 BP cases and 258,900 controls, giving a prevalence of BP of 26%. Cases and controls did not differ significantly by age (mean age 57.05 years) or sex (54% female) (Supplementary Table 1). SNP-based heritability estimated by LD score regression from the BP GWAS was 4% on the observed scale and 7.3% on the liability scale. LD-score regression estimated the genomic inflation factor to be 1.29 with intercept of 1.032±0.009 (giving the estimate of standardized genomic control inflation factor of λ_1000_=1.00024), suggesting that most of the inflation was introduced by polygenic effects and that the influence of confounding by population structure and cryptic relatedness was minimal (QQ-plot in Supplementary Figure 1).

After adjusting the results of the discovery GWAS for genomic control factor of 1.032, a total of 183 SNPs positioned over 5 loci remained statistically significant at genome-wide significance level of p≤5×10^−8^ (Figure 2; Table 1). Conditional and joint analysis confirmed that all 5 regions were independent of one another (Supplementary Table 2A). Using meta-analysis for the UK Biobank replication cohorts and the CHARGE Consortium cohorts (total N = 154,970-157,752), three associations were replicated (p<0.01) (Supplementary Table 2B): rs12310519 (p = 5.00×10^−5^), rs7814941 (p = 5.32×10^−5^), and rs3180 (p = 6.59×10^−3^).

**Table 1.**
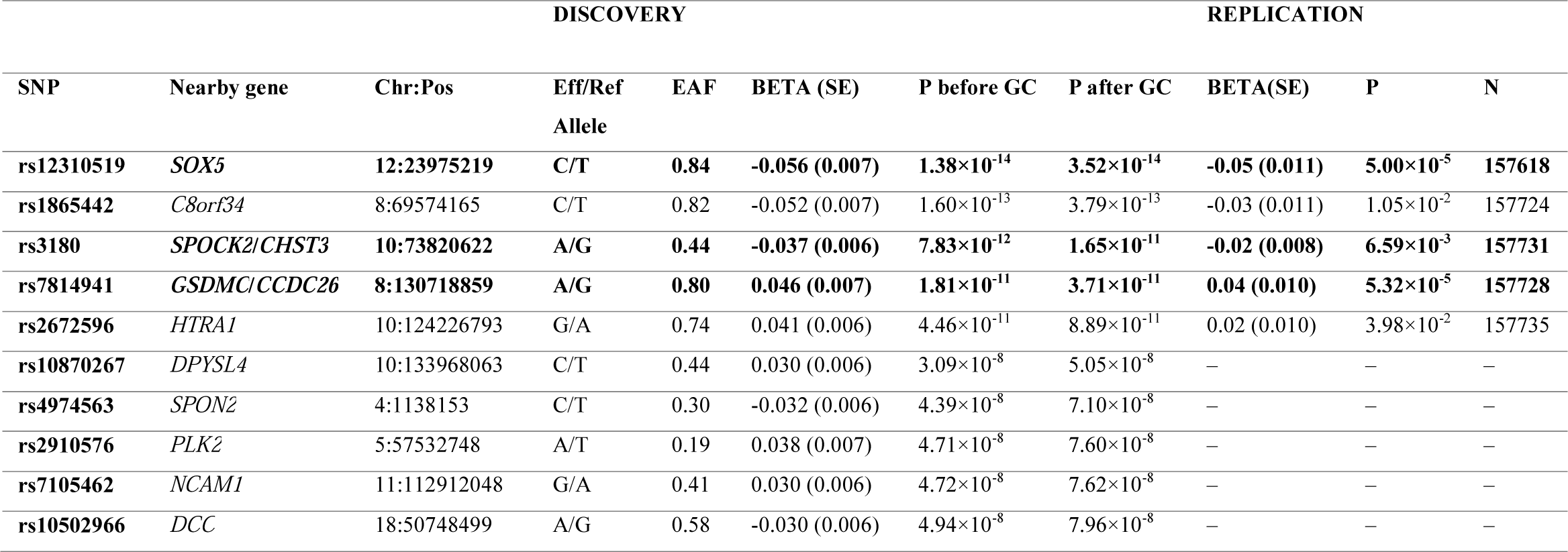
Results of discovery and replication GWAS of BP. **Table legend:** Chr:Pos – physical position of the SNP; Eff/Ref Allele – Effect and Reference alleles; EAF – effect allele frequency; BETA (SE) – effect of the SNP and standard error; P before GC – p-value before genomic control; P after GC – p-value after genomic control; N – sample size of replication. Bold font indicates the SNPs that passed the thresholds for statistical significance (5×10^−8^ for the discovery and 0.01 for replication phases, respectively).

**Figure 2.**
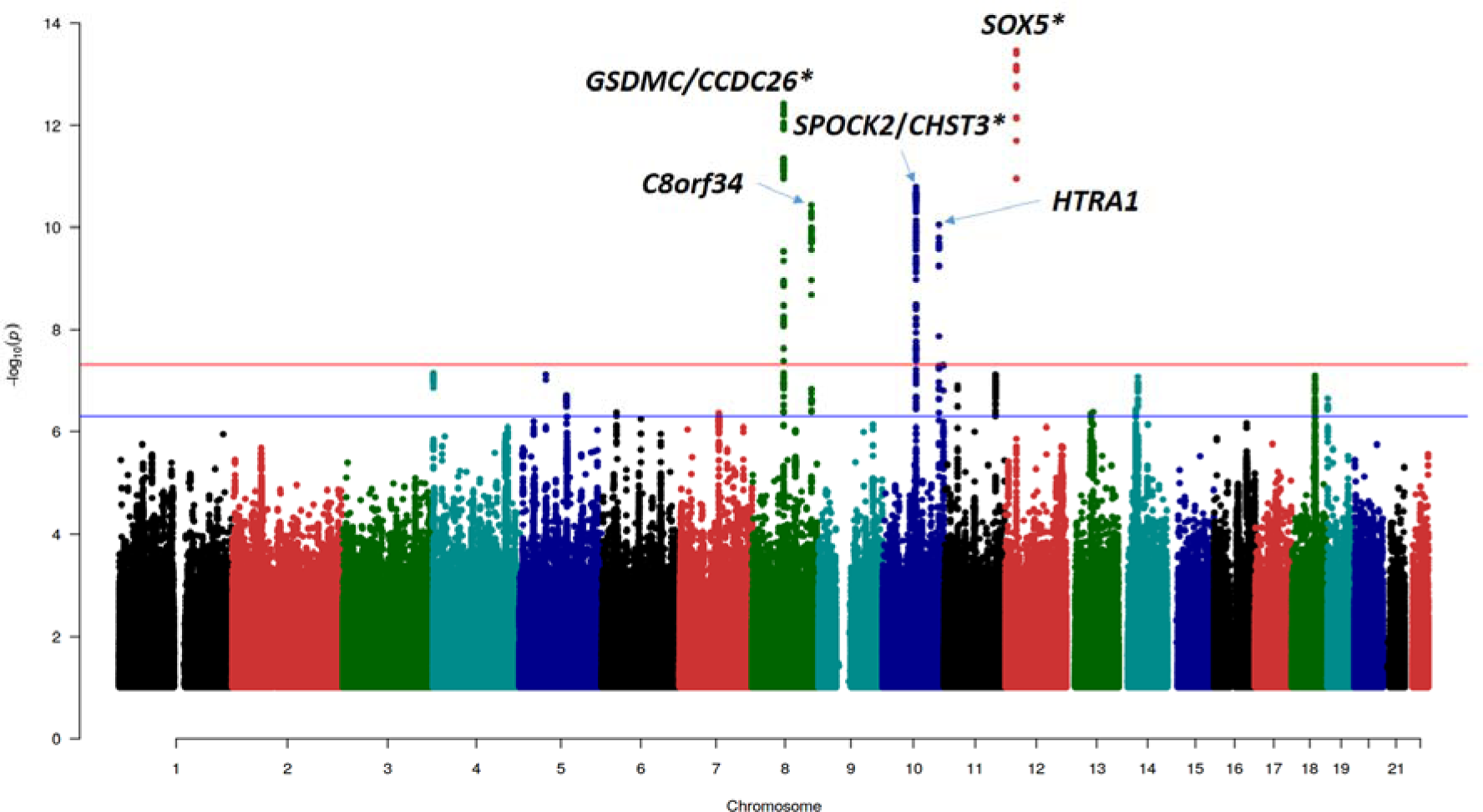
Manhattan plot of discovery GWAS for back pain. Correction was made for genomic control (1.032). The red line corresponds to genome-wide significance threshold of 5×10^−8^, while the blue line corresponds to a suggestive association threshold of 5×10^−7^. Only SNPs with p<0.1 are presented. Asterisks depict replicated loci.

Of the three replicated loci, two have been reported previously as associated with other BP phenotypes: the chromosome 12 lead SNP rs12310519 located in the intron of the *SOX5* gene was associated with chronic BP in the recent GWAS by the CHARGE and PainOMICS consortia ^22^. The region on chromosome 8 (lead SNP rs7814941), located in an intergenic site of *GSDMC* and *CCDC26* was identified in a study of sciatica,^23^ and was among the loci associated with chronic BP at p<5×10^−8^ in the GWAS by the CHARGE and PainOMICS consortia,^22^ but not previously replicated.

The novel replicated locus on chromosome 10 (rs3180 SNP) lies in the region between the 3’-UTR of *SPOCK2* and downstream of *CHST3* genes. This region was previously shown to be associated with LDD with the leading SNP rs4148941 reported as a functional variant influencing the *CHST3* gene expression level in intervertebral disc tissue ^24^. This gene encodes an enzyme which catalyzes sulfation of chondroitin, a component of proteoglycans crucially important in cartilage tissue function and hydration. Rare mutations in *CHST3* that disrupt its enzymatic activity have been reported in patients with recessive skeletal abnormalities, including spondyloepiphyseal dysplasia Omani type, Larsen syndrome, humero-spinal dysostosis, and chondrodysplasia with multiple dislocation ^25–29^. Another gene in the region, *SPOCK2*, was previously reported as the positional candidate for bronchopulmonary dysplasia ^30^, chromosome 16q carcinogenic deletion (along with *CHST3*) ^31^, and age of smoking initiation ^32^. The gene encodes a proteoglycan SPARC/Osteonectin (Cwcv And Kazal Like Domains Proteoglycan 2) involved in extracellular matrix formation and is highly expressed in the central nervous system (CNS) ^33^. Using available *in-silico* instruments we did not find sufficient evidence to determine whether *SPOCK2* or *CHST3* was the most likely gene associated with BP on chromosome 10 (See Additional results file).

To achieve higher statistical power for the subsequent study of pleiotropic effects and genetic correlations, a meta-analysis of discovery (EA British N = 350,000) and replication sets (other EA N = 103,862) was performed, giving total N = 453,862. The SNP-based heritability estimate from this GWAS was 4.2% on the observed scale and 7.7% of the liability scale. LD-score regression estimated the genomic inflation factor to be 1.37 with intercept of 1.036±0.009 (standardized genomic control inflation factor of _1000_=1.00021). A total of 651 SNPs at 23 loci achieved genome-wide significance threshold of p<5×10^−8^ (Supplementary Figure 2, Supplementary Tables 3). COJO analysis confirmed that significant loci were all independent of each other. Subsequently, we refer to the results of this meta-analysis as BP_ma_ to contrast with the discovery GWAS.

### Causal and pleiotropic effects of genetic factors underlying back pain and its risk factors

#### Identifying causal genes via a study of gene expression

For replicated regions we aimed to identify genes whose expression might mediate the association between SNP and BP. We performed a summary-data based Mendelian randomization (SMR) analysis followed by heterogeneity in dependent instruments (HEIDI) analysis ^34^ using eQTL data from a range of tissues including blood ^35^ and 44 tissues provided in the GTEx v. 6p database ^36^ (Supplementary table 4A). In short, SMR tests the association between gene expression in a particular tissue and a trait using the most highly associated SNP as a genetic instrument. A significant SMR test indicates that a given functional variant determines both gene expression and the trait of interest via causality or pleiotropy, but it may also suggest that functional variants underlying gene expression are in linkage disequilibrium with those controlling the trait. Inference of whether a functional variant mediates both BP and gene expression were made based on the HEIDI test: P_HEIDI_ > 0.01 (likely shared causal SNP) and P_HEIDI_ < 0.01 (sharing of a causal SNP is unlikely). Results are presented in Supplementary Table 4B.

We observed a statistically significant SMR (p<3×10^−5^) and no difference in association patterns for the rs3180 locus and *SPOCK2* in blood (β_SMR_ = 5.9; p_SMR_ = 1.0×10^−8^) and in adrenal gland (β_SMR_ = −20.6; p_SMR_ = 1.3×10^−6^). Moreover, for this locus we detected three probes with suggestive significance SMR coefficient and p_HEIDI_ >0.01: two probes for *CHST3* gene in testis (β_SMR_ = 6.3; p_SMR_ = 5.5×10^−5^) and in EBV-transformed lymphocytes (β_SMR_ = −17.1; p_SMR_ = 1.7×10^−4^); and one probe for *SPOCK2* gene in muscle skeletal tissue (β_SMR_ = −9.6; p_SMR_ = 8.7×10^−5^). The results suggest that either *SPOCK2* or *CHST3* or both are possible causal genes for BP in the region tagged by rs3180. It is worth noting, though, that some of the tissues with significant findings in this analysis (testis, EBV-transformed lymphocytes, adrenal gland) do not seem relevant to BP in an anatomical or functional sense. Nevertheless, a BP-associated variation in *SPOCK2*/*CHST3* region was linked with *CHST3* expression in intervertebral disc tissue in an *in vitro* functional study previously ^24^.

For the locus tagged by rs7814941, we detected two expression probes with statistically significant SMR coefficients that corresponded to *GSDMC* gene. However, in all cases there was a significant (p_HEIDI_ <2×10^−7^) difference in association patterns between the SNP and gene expression and BP. This suggests that the association between this region and BP is unlikely driven by variation in *GSDMC* gene expression.

#### Pleiotropic effects of genetic variants associated with BP and other complex traits

Using the SMR/HEIDI approach, we also tested for potential pleiotropy of effects of three BP loci on seventeen known risk factors or related conditions for which data were available in public databases: osteoarthritis, self-reported intervertebral disc problems, osteoporosis, scoliosis, smoking status, standing height, BMI, well-being (happiness), fluid intelligence score, educational attainment (years of education), anxiety/panic attacks, depression and the ‘Big Five’ personality traits (openness to experience, conscientiousness, extraversion, agreeableness, and neuroticism) (Supplementary Table 4C; Supplementary methods).

Results are presented in Supplementary Table 4D. Statistically significant (p<9.8×10^−4^) SMR coefficients were revealed for height with variants rs7814941 and rs3180 (p_SMR_ = 3.60×10^−13^ and 4.35×10^−5^, respectively). Locus rs7814941 showed significant heterogeneity in association patterns with height (p_HEIDI_ = 5.58×10^−12^) suggesting the presence of different functional variants for height and BP at this locus. Locus rs3180 showed no heterogeneity in association patterns between BP and height in HEIDI (p_HEIDI_ = 0.79), thus suggesting pleiotropy – the same functional variant(s) was influencing both traits. All three loci demonstrated significant SMR results with intervertebral disc problems (p_SMR_ = 3.30×10^−7^, 3.75×10^−7^, and 3.17×10^−5^, for rs3180, rs12310519, and rs7814941, respectively; Table 2); with all three showing no heterogeneity in association patterns between BP and intervertebral disc problems (all p_HEIDI_ > 0.01); and in all cases SMR coefficient was positive suggesting that the same causal genetic factors attributable to these loci increase the risk of both BP and self-reported intervertebral disc problems.

**Table 2.** Results of summary-level Mendelian randomization and pleiotropy analysis of SNPs associated with BP. **Table legend:** Results of SMR/HEIDI tests using data from GeneAtlas. For the HEIDI tests, a hypothesis of pleiotropy was rejected at p < 0.01; with p > 0.01, we considered pleiotropy as a likely explanation. Two traits (height and intervertebral disc problems) with at least one significant SMR coefficient (p<9.8×10^−4^) among three loci are presented. β_SMR_ is SMR coefficient; p_SMR_ is p-value for SMR test; p_HEIDI_ is p-value for HEIDI test (not calculated if p_SMR_ was insignificant, “–”).

| Trait | Statistics | SNP (Gene) |  |  |
| --- | --- | --- | --- | --- |
|  |  | rs3180<br>(*SPOCK2/CHST3*) | rs12310519<br>(*SOX5*) | rs7814941<br>(*GSDMC/CCDC26*) |
| Height | $\beta_{\text{SMR}}$ | -1.677 | 0.800 | 6.013 |
| | $p_{\text{SMR}}$ | $4.3 \times 10^{-5}$ | $7.4 \times 10^{-3}$ | $3.6 \times 10^{-13}$ |
| | $p_{\text{HEIDI}}$ | 0.79 | – | $5.6 \times 10^{-12}$ |
| Intervertebral<br>disc problems | $\beta_{\text{SMR}}$ | 0.068 | 0.050 | 0.040 |
| | $p_{\text{SMR}}$ | $3.3 \times 10^{-7}$ | $3.8 \times 10^{-7}$ | $3.2 \times 10^{-5}$ |
| | $p_{\text{HEIDI}}$ | 0.75 | 0.12 | 0.50 |

### Back pain shares genetic components with psychiatric, sociodemographic and anthropometric traits

To establish shared genetic components between BP and other complex traits, we carried out an agnostic analysis of 225 complex traits available in LD-hub. We observed a significant genetic correlation (p<4.4×10^−5^) between BP_ma_ and 33 traits (Supplementary Table 6, Supplementary Figure 3), with the strongest positive correlations (ρ_g_>0.35) found with BP and neuroticism ^37^ (ρ_g_=0.49), insomnia ^38^ (ρ_g_=0.46), depressive symptoms ^37^ (ρ_g_=0.53) and major depressive disorder ^39^ (ρ_g_=0.39). The strongest negative correlations (ρ_g_<−0.35) were between BP_ma_ and age of first birth ^40^ (ρ_g_=−0.49), years of schooling ^41^ (ρ_g_=−0.47), mothers age at death ^42^ (ρ_g_=−0.43), parents age at death ^42^ (ρ_g_=−0.38) and college completion ^43^ (ρ_g_=−0.51). The traits exhibiting strong genetic correlation with BP fell into several distinct clusters (Figure 3): 1) the cluster of obesity-related traits, 2) the cluster related to mood and sleep, and 3) the cluster related to sociodemographic factors (including education) and smoking.

**Figure 3.**
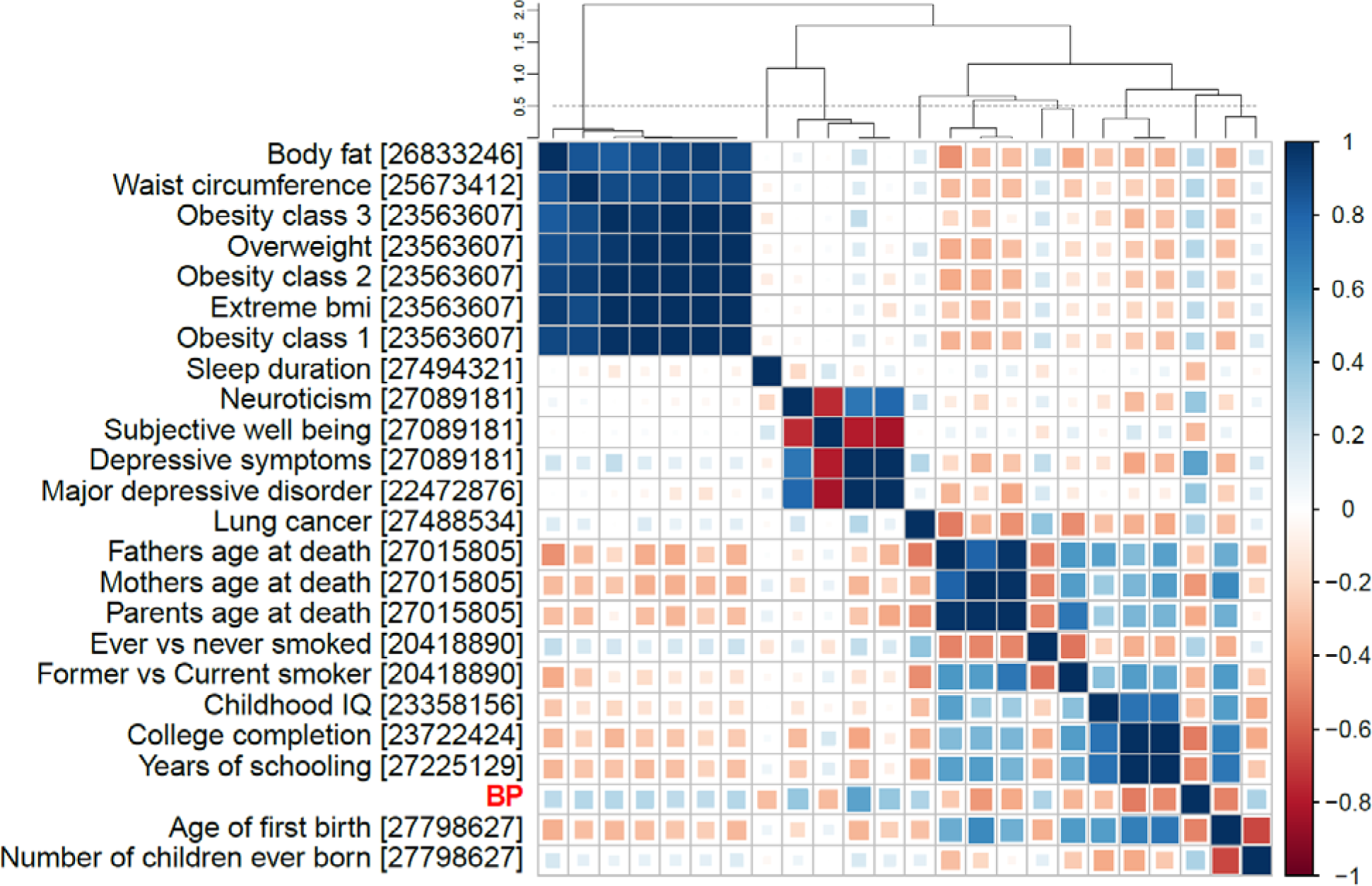
Heatmap for 23 traits with strongest statistically significant genetic correlations with back pain (absolute ρ_g_ ≥ 0.25; p≤4.4×10^−5^). Hierarchical clustering was carried out based on genetic correlations between all pairs of traits. PMID references are placed in square brackets. The dashed line on the cluster dendrogram refers to the threshold of 0.5, depicting 9 subclusters (including BP).

To identify which pair-wise genetic correlations were conditionally independent of each other, we calculated partial genetic correlations for BP_ma_ and 8 traits selected from each subcluster (using a distance threshold of 0.5 on a hierarchical clustering dendrogram) of the genetic correlation matrix (Figure 4, Supplementary Figure 4). In short, partial correlation is the measure of association between two variables while controlling for the effect of one or more additional variables. This analysis found such traits as “mother age of death”, “lung cancer”, and “former vs current smoking”, and “age of first birth” to not be independently correlated with BP. Partial correlations for depressive symptoms and sleep duration were similar to the pair-wise correlations. Finally, partial correlations with BP for “waist circumference” and “college completion” were much smaller than the pair-wise correlations but remained statistically significant (p<4.4×10^−5^).

**Figure 4.**
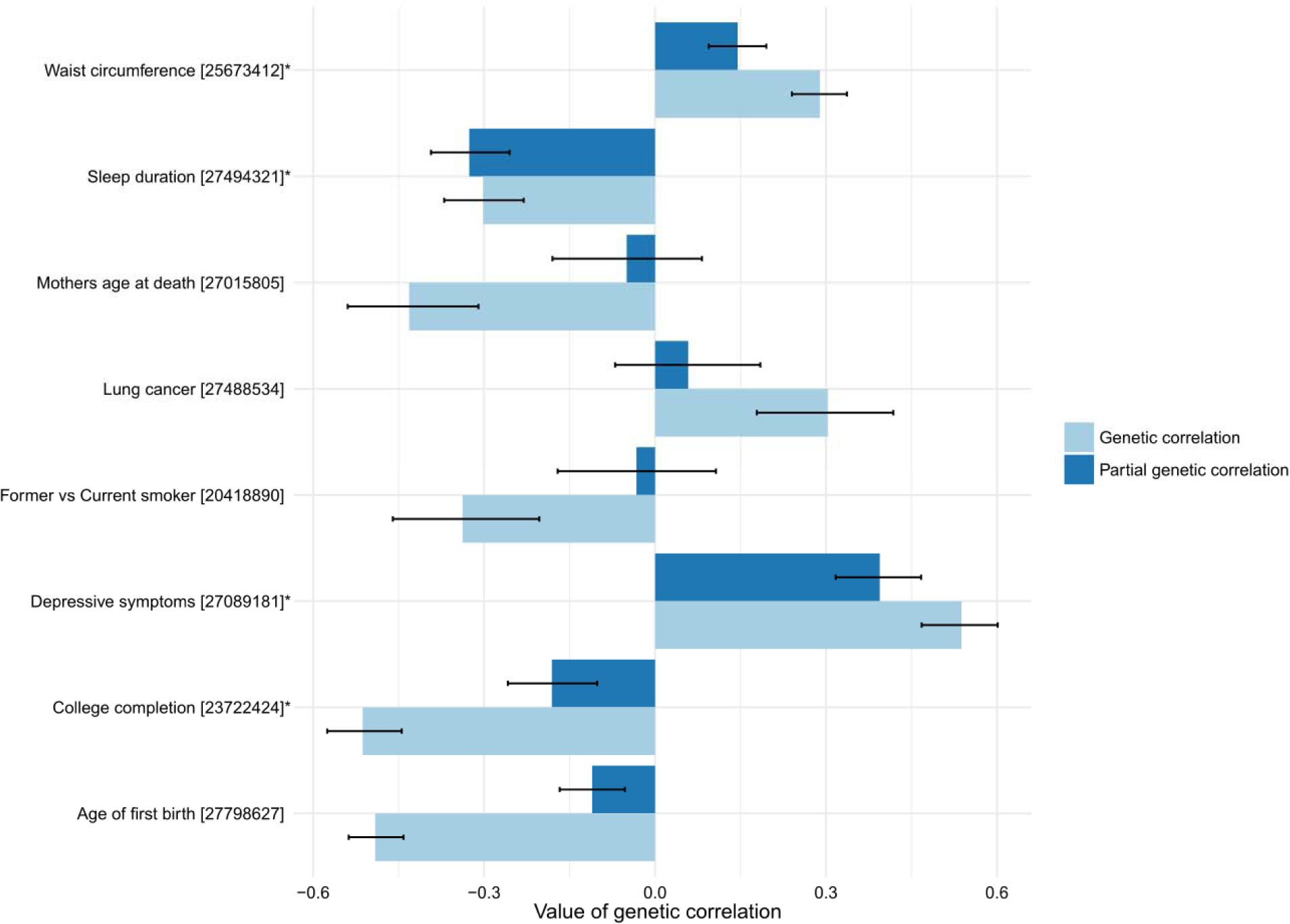
Partial genetic correlation and pair-wise genetic correlation barplots for 8 traits (one trait from each subcluster with threshold of 0.5 on hierarchical clustering dendrogram of genetic correlation matrix). Error bars correspond to 95% confidence intervals. Asterisks depict traits in which partial correlation with BP is significant (p<4.4×10^−5^).

In addition to an agnostic analysis of all complex traits from LD-hub, we also carried out a focused analysis of genetic correlations between BP and 17 complex traits considered as risk factors for BP: self-reported osteoarthritis, self-reported intervertebral disc problems, self-reported osteoporosis, scoliosis, smoking status, standing height, BMI, happiness, fluid intelligence score, years of education, anxiety/panic attacks, depression and Big Five traits (Supplementary Table 4C). The strongest positive correlations were found for self-reported intervertebral disc problems (ρ_g_ = 0.77, p = 6.7×10^−24^); self-reported osteoarthritis (ρ_g_ = 0.55, p = 7.5×10^−41^); and depression (ρ_g_ = 0.44, p = 1.3×10^−23^). The strongest negative correlation was found for education attainment (ρ_g_ = −0.47, p = 7.1×10^−101^). Scoliosis, smoking status and BMI had moderate positive genetic correlation with BP_ma_ (ρ_g_ = 0.35, 0.35 and 0.33 respectively, with p=0.001, 7.3×10^−42^ and 2.0×10^−56^ respectively). Overall, the results of the analysis of the risk factors were consistent with the analysis of 225 traits.

### Genetic factors underlying back pain are involved in neurological pathways

We used DEPICT with all independent variants (as identified by COJO analysis) from BP_ma_ with p < 1e-5 (227 SNPs in total; Supplementary Table 6A-C) and identified potential enrichment of gene sets (FDR<0.2) related to nervous system development and skeletal muscle development. We did not identify a significant enrichment of expression across any tissues and cell types (FDR > 0.2), although we observed a trend towards enrichment of components of CNS (Supplementary Table 7A-C). Similar results were observed when analyzing enrichment of expression of genes located around 23 BP_ma_ independent genome-wide significant variants (Supplementary Table 7D-F).

Analysis by MAGMA ^44^ revealed three significant gene sets (Supplementary Table 8): M12307 (“Nikolsky breast cancer 16q24 amplicon”, FDR=0.02; copy number amplicons of 53 genes enriched with major tumorigenic pathways and breast cancer-causative genes ^45^), GO:0051590 (positive regulation of neurotransmitter transport, FDR=0.02) and GO:0021952 (central nervous system projection neuron axonogenesis, FDR=0.02). Tissue expression analysis for 30 general tissue types revealed significant enrichment of expression in brain (FDR=0.01).

## DISCUSSION

The current study is the largest genetic association study to date for BP and included more than 500,000 individuals. The results provide insights into the genetic composition of predisposition to BP, one of the leading causes of disability worldwide. We quadrupled the number of genome-wide significantly associated BP loci (from five ^22^ to 23), and increased the number of replicated BP loci from one to three. Our work has implicated two new positional candidate genes: *SPOCK2* and *CHST3*. The region where these genes reside has previously been described as associated with LDD in Chinese individuals, and *in vitro* functional study suggested a mechanism linking variation in this locus (specifically, rs4148941) and expression of *CHST3*, a functionally highly plausible gene ^24^. Our *in silico* functional analysis, however, suggests that the closely adjacent *SPOCK2* gene may be another candidate in the region. In particular, we provide evidence of relationships between both *SPOCK2* and *CHST3* gene expression and the risk of BP. At the same time, using available *in-silico* instruments, we couldn’t provide enough evidence in favor of *SPOCK2* or *CHST3* as the most likely gene associated with BP on the locus on chromosome 10 (see Additional results file).

We found evidence of pleiotropic effects for the genetic factors underlying BP, height, and intervertebral disc problems. From epidemiological studies, both height and LDD are known to be associated with BP and have been proposed to have causal effects on BP ^5,46^. The genetic pleiotropy identified in the current study provides insight into the molecular background underlying these associations. Importantly, while only one of the three loci (rs3180) exhibited pleiotropic effects for BP and height, all three loci demonstrated pleiotropy for BP and self-reported intervertebral disk problems. In addition, the observed genetic correlation between height and BP was small and statistically insignificant (ρ_g_=0.05, p=0.07), while the genetic correlation between intervertebral disc problems and BP was high and statistically significant (ρ_g_ = 0.77, p = 6.7×10^−24^). These results suggest stronger shared underlying genetic factors between intervertebral disk degeneration and BP, as compared to height and BP, in keeping with the epidemiological evidence of a stronger association of BP with disc degeneration ^16,47^ and a weaker association with height.^48^ An alternative explanation for our observation that loci influencing BP also affect intervertebral disc problems might be an overlap between individuals reporting BP and intervertebral disc problems in UK Biobank. However, at least for the lead SNPs tagging regions near *SOX5* (rs12310519), and *GSDMC/CCDC26* (rs7814941 via proxy rs4733724), nominally significant associations (p = 1.1×10^−4^ and p = 0.023, respectively) with MRI-proven LDD were found in a meta-analysis of 4600 individuals independent from the current study sample and not selected by BP status.^49^ This strengthens the argument in support of a shared genetic basis for BP and intervertebral disc problems.

To our knowledge, this is the first study to use contemporary quantitative genetic methods in an attempt to replicate the results of twin studies examining shared genetic influences on BP with other traits, including putative BP risk factors.^10,16,47,50–54^ In so doing, we took the broadest approach to date and examined a wide range of complex traits and known risk factors, revealing three clusters sharing significant genetic correlations with BP : the obesity-related traits, the mood and sleep related traits, and the sociodemographic factors (including education) and smoking. Moreover, we identified mutually independent genetic correlations between BP and depression, sleep disturbance, waist circumference and college completion. The magnitude and direction of many of the observed genetic correlations in the current study follow from the results of classic epidemiology and genetic epidemiology studies of BP suggesting, perhaps, that the environmental components to these risk factors have been overstated or at least themselves have a genetic basis. For instance, we observed strong positive genetic correlations between BP and depression related phenotypes, and between BP and obesity-related traits. These traits are known to often co-occur with BP and twin studies have suggested that they share underlying genetic factors^17^, with similar genetic correlations also seen for other pain phenotypes.^55–57^ Our results confirm a recent twin report of genetic correlation of sleep disturbance with BP.^58^ Overall, the analysis of genetic correlations provides evidence for shared molecular pathways underlying BP and traits considered as BP risk factors, thus providing basis for identification of causal links between them.

Our pathway analysis revealed the importance of genetic factors in CNS and skeletal muscle in BP. While the CNS has long been recognized as the key component in the pathogenesis of chronic pain, ^59^ the role of skeletal muscle is still not well defined.^60,61^ Altogether, these data provide a starting point for further functional analyses of mechanisms underlying BP (Figure 5). The study of pleiotropy and genetic correlations, supported by the pathway analysis, suggests at least two strong molecular axes of BP genesis, one related to structural/anatomic factors such as intervertebral disk problems and anthropometrics; and another related to the psychological component of pain perception and pain processing. These two axes correspond roughly to the different “biomedical” and “biopsychosocial” viewpoints that have dominated BP research and clinical care for the past several decades.^62^ Pathway analysis also produced an unexpected enrichment for genes involved in “Nikolsky breast cancer 16q24 amplicon” gene set. This gene set includes 53 genes and represents one of 30 genomic regions with copy number gain found in the analysis of 191 breast tumours ^45^. It is not known to be enriched for pain-related or other relevant pathways; therefore, its relationship with BP needs to be explored further.

**Figure 5.**
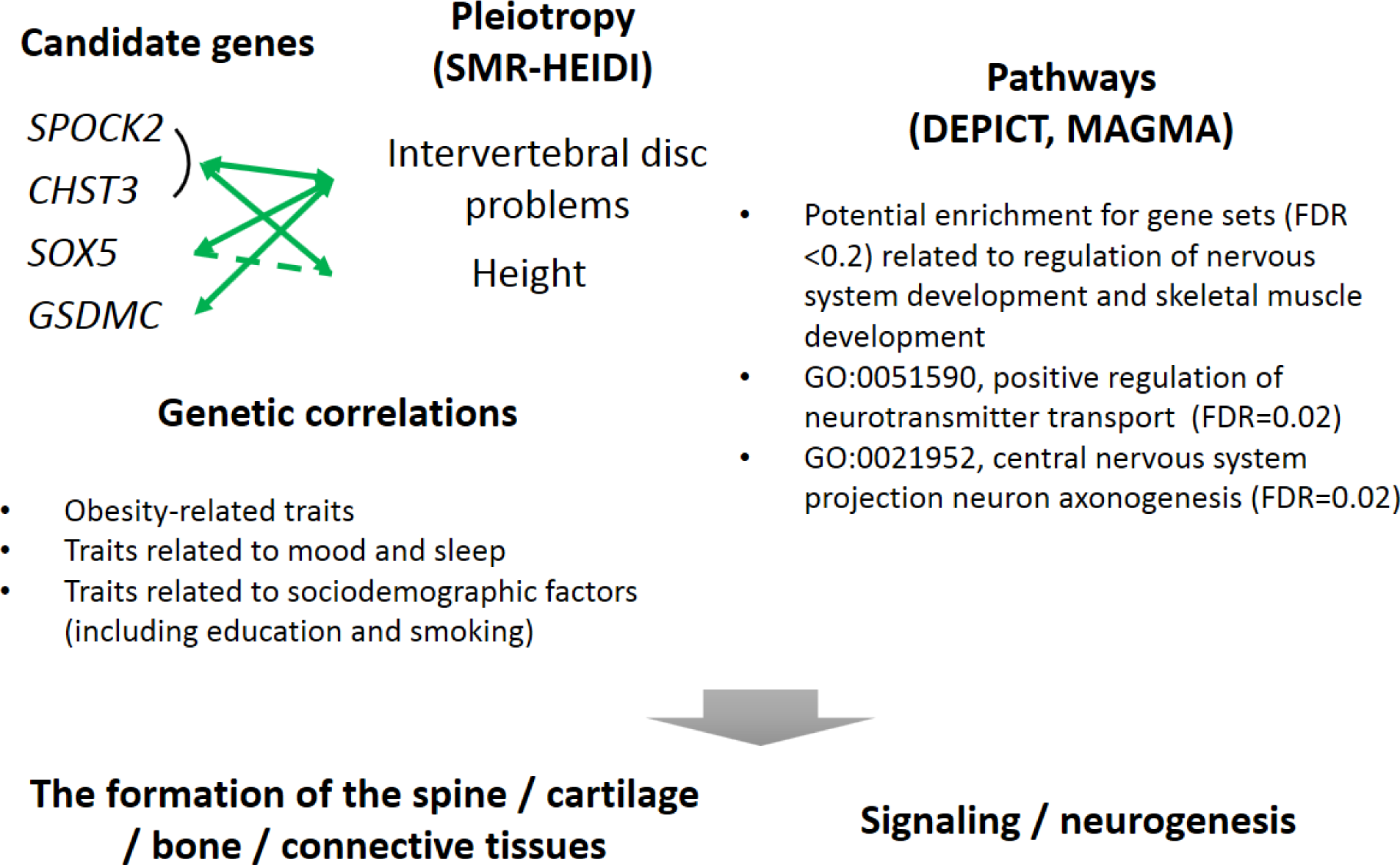
Summary of genes and pathways in back pain. Left part of the figure summarizes information about positional candidate genes and genetic correlations. Green arrows depict pleiotropy by SMR/HEIDI method. Dashed green lines depict suggested pleiotropy by SMR/HEIDI. Left part of the figure summarizes the results of pathway analyses.

Despite the study of close to half a million people, we identified and replicated only 3 loci. Together with the evidence of relatively high heritability from twin studies ^10–13^, this suggests that BP is genetically a very complex, highly polygenic phenotype. In part, this can be explained by the heterogeneity of the phenotype itself, as BP likely occurs for a vast range of reasons, many having different underlying molecular pathology^63^. Our approach used a standard definition of “any back pain” in the discovery stage, but permitted some heterogeneity with respect to BP duration among cohorts included in the replication stage (any BP in the UK Biobank sub-cohorts vs chronic BP in the CHARGE cohorts). Future progress of genetic studies of BP would benefit from more consistent phenotyping. On the other hand, our experience to date of working with back pain consortia – with no more than 3 cohorts having closely comparable back pain definitions and question items – has shown that it is extremely difficult to bring together cohorts of comparable size having uniform phenotype definition. This reflects the current state of BP research, where there is no universally accepted gold standard for defining BP.^22^ Recently established consensus guidelines for core BP definitions may facilitate future efforts to harmonize definitions between cohorts.^64^

## ONLINE METHODS

### Phenotype definition

The study was based on data from the UK Biobank Resource ^21^ and cohorts from CHARGE Consortium Musculoskeletal Working Group. For UK Biobank cases of BP were defined as those who reported “Back pain” in the response to the question: “Pain type(s) experienced in last month”. Controls were defined as those who did not report BP in response to this question. Individuals who did not reply or replied: “Prefer not to answer” or “Pain all over the body” were excluded.

For CHARGE Consortium cases were defined as those reporting BP present for at least 3 months, while the controls were defined as those who reported no BP or BP with shorter duration.^65^ Thus, the definition of BP in these cohorts corresponded to chronic BP.

### Sample

The available sample from UK Biobank included 487,409 individuals with imputed data. For the discovery set we selected at random 350,000 British individuals of European ancestry (EA) according to the genetic principal components provided by the UK Biobank (Supplementary Table 1).

For replication, we used a combination of the UK Biobank participants not included in the discovery set, and from the CHARGE Consortium ^22^. Replication cohorts from the UK Biobank comprised rest of EA individuals (n = 103,862), individuals of African ancestry (AA, n = 7,259), individuals of South Asian ancestry (Indian, Pakistani, and Bangladeshi; n = 7,159), and Chinese individuals (n = 1,485). The CHARGE Consortium provided data for EA individuals from 15 cohorts (total n = 35,205-37,987). To reduce the risk of bias due to population stratification, all these groups were analysed separately followed by a meta-analysis. Total resulting sample size for replication was 154,970-157,752 individuals (Supplementary Table 1).

### Statistical analysis

#### Genome-wide association testing

PLINK 2.0 was used to carry out the genome-wide association analysis in the UK Biobank discovery and replication samples. Imputed genotype provided by the UK Biobank were used ^21^ and only SNPs from the list of HRC imputed SNPs were analysed (http://www.haplotype-reference-consortium.org/site) due to reported issue with SNPs imputed using 1000 Genomes panel (http://www.ukbiobank.ac.uk/2017/07/important-note-about-imputed-genetics-data/). Logistic regression was used to evaluate additive genetic effects of the single nucleotide polymorphisms (SNPs) for BP as a binary trait adjusting for age, sex, genotyping array type, and 10 genetic principal components provided by UK Biobank.

The following filters were applied: minor allele count ≥100; deviation from Hardy-Weinberg equilibrium p-value ≥1e-6; genotyping call rate ≥0.98%; individual call rate ≥0.98%; and imputation quality score ≥0.7 (MACH r2 calculated by PLINK 2.0). Only biallelic markers were used and SNPs that had the same rsID in different genomic locations were excluded.

#### Conditional and joint multi-SNP analysis

Conditional and joint analysis (COJO) as implemented in the program GCTA ^66^ was used to find SNPs independently associated with the phenotype. As the input, this method uses summary statistics and a reference sample that is utilised for the LD estimation. We performed the analyses using p = 5×10^−8^ and p = 1×10^−5^ as the genome-wide significance and suggestive significance thresholds respectively. For the LD reference, we used a sample of 10,000 British EA individuals randomly selected from 350,000 people used in the GWAS discovery phase.

#### Replication and meta-analysis

Replication was performed by meta-analysis of all replication cohorts for loci selected at the discovery phase. Replication significance threshold was set as p-value<0.01 (Bonferroni corrected 0.05/5). Subsequent analyses of heritability, genetic correlation, and functional investigation used the results of meta-analysis of the discovery cohort and replication cohort of EA individuals from UK Biobank (N=453,862). METAL software ^67^ was applied using inverse-variance-weighted approach.

LD hub ^68^ and ldsc ^69^ tools were used to estimate SNP-captured heritability. Summary statistics files were filtered using ldsc software with default options (r2>0.9, MAF>0.01 and the overlap with “high quality SNPs” – a total of 1,215,001 common HapMap3 SNPs with high imputation quality). The HLA region on chromosome 6 was excluded. These SNPs were used for the further analysis of heritability and genetic correlations as well as to estimate genomic control inflation factor lambda (intercept) ^70^.

### Genetic correlation analyses

Genetic correlations were estimated using the BP meta-analysis results (N=453,862), not including the CHARGE cohorts. Two sets of traits were analyzed. First included a total of 225 traits available on LD-hub after removing duplicates via using only the most recent study for each trait as indicated by the largest PMID number. Another set comprised traits considered by us as risk factors for BP: self-reported osteoarthritis, self-reported intervertebral disc problems, self-reported osteoporosis, scoliosis, smoking status, standing height, BMI, happiness, fluid intelligence score, years of education, anxiety/panic attacks, depression and Big Five personality traits (openness to experience, conscientiousness, extraversion, agreeableness, and neuroticism). Genetic correlations between BP and 225 traits were considered statistically significant at p-value ≤ 4.4×10^−5^ (Bonferroni corrected, 0.01/225). To visualize the results, we focused on genetic correlations of greatest magnitude and selected only the traits with absolute values of genetic correlation with BP >0.25. This filtering led to a total of 23 traits (excluding BP). Clustering and visualization was carried out using “corrplot” package for R and basic “hclust” function. For clustering, we estimated squared Euclidean distances by subtracting absolute values of genetic correlation from 1 and used Ward’s clustering method.

To obtain genetic correlations that were independent from each other, we estimated partial genetic correlations for a subset of traits using the inverse of correlation matrix (free of collinearity) followed by the correlation estimate given by the equation 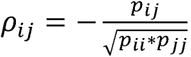. We selected for this analysis one trait from each subcluster (using distance threshold of 0.5 on hierarchical clustering dendrogram) of the matrix of genetic correlations for 24 traits (we selected only the trait with the highest absolute value of its correlation with BP). This selection led to 8 traits in total.

### *In silico* functional analysis

#### VEP and credible set

Functional annotation of SNPs was carried out using variant effect predictor (VEP) software ^71^ with GRCH37 genomic reference. For each studied locus we selected the set of SNPs (credible set) that had strong associations and the most probably had causal influence on BP. We used PAINTOR software ^72^ to prepare the credible set of SNPs. For this analysis, we provided PAINTOR with clumping results, LD matrices and annotation files. Using PLINK1.9 and 10,000 samples reference set described above (the same subset as used in COJO and DEPICT analyses) we performed clumping analysis with ‘p1’ and ‘p2’ p-value threshold parameters set to 5×10^−8^, ‘r2’ set to 0.1 and MAF>0.002. Then, using the same reference set we generated pair-wise correlation matrix for all SNPs in each region in clumping analysis results using PLINK “--r” option. When running PAINTOR, we did not use annotations; we changed options controlling input and output files format only, and otherwise we have used default parameters. In the next step, all output results were aggregated into one file and SNPs marked by PAINTOR as 99% credible set were chosen for further functional annotation.

#### SMR/HEIDI analysis

Potential pleiotropic effects of genetic variants on BP and other traits were tested using summary data-based Mendelian randomization (SMR) analysis and heterogeneity in dependent instruments (HEIDI) method ^34^. SMR-HEIDI is analogous to conventional Mendelian randomization and may be conducted using summary level GWAS data. In short, the first method tests for association between the traits of interest mediated by a locus, and the second test identifies whether the traits are affected by the same underlying causal variant. This analysis was carried out for SNPs associated with BP in the current study. Briefly, starting with an index SNP, we screened for traits which may be affected by genetic variation in the same region, and then performed a pleiotropy vs linkage disequilibrium test. In the screening stage, we used a limited list of traits including 19 traits considered risk factors for BP (19 traits in total; Supplementary table 4C). To perform HEIDI analysis, regional summary level GWAS results are required, including regression coefficients and respective standard errors. Such data were available for seventeen traits: self-reported osteoarthritis, self-reported intervertebral disc problems, self-reported osteoporosis, scoliosis, smoking status, standing height, BMI, happiness, fluid intelligence score, years of education, anxiety/panic attacks, depression and Big Five personality traits (openness to experience, conscientiousness, extraversion, agreeableness, and neuroticism) (Supplementary table 4C).

We also examined for overlap between the SNPs associated with BP and eQTLs in blood ^35^ and 44 tissues provided by the GTEx database ^36^ (Supplementary tables 4A) using a similar procedure: we examined if a SNP was available for specific expression probe in the region of interest and, if positive, the HEIDI test was performed as above.

Following Bonferroni procedure, the results of the SMR test were considered statistically significant at p < 3×10^−5^ (0.05/1685, where 1685 is the number of probes available in blood eQTL data and GTEx data for three studied loci) for eQTLs; and p<9.8×10^−4^ (0.05/(17×3) accounting for 3 studied loci and 17 complex traits) for complex traits.

For the HEIDI tests, a hypothesis of pleiotropy was rejected at p < 0.01; the hypothesis was accepted at p>0.01.

#### Gene prioritization, pathway and tissue enrichment analysis

To prioritize genes in associated regions, gene set enrichment and tissue/cell type enrichment analyses were carried out using DEPICT software v. 1 rel. 194 ^73^. For this analysis we chose independent (by COJO) variants found in the BP meta-analysis results (N=453,862) with p<5×10^−8^ (23 SNPs) and p<1×10^−5^ (227 SNPs). We used a random subset of 10,000 individuals from the UK Biobank for calculation of LD (the same subsets as used for COJO analysis).

We also conducted gene analysis and gene-set analysis using MAGMA v1.6 included in FUMA web tool ^44^. Reference genome was 1000 genomes phase 3. The MAGMA output from FUMA was based on summary statistics. We used standard options suggested by FUMA web tool for this analysis.

## SUPPLEMENTARY MATERIALS LEGEND

**Supplementary Figure 1 –** QQ plot for discovery GWAS.

**Supplementary Figure 2 –** Manhattan plot of meta-analysis GWAS of discovery and replication samples from the UK Biobank (N=453,862). Red line corresponds to the genome-wide significance threshold of 5×10^−8^, while blue line corresponds to the suggestive association threshold of 5×10^−7^.

**Supplementary Figure 3 –** Barplots of genetic correlations between back pain and complex traits showing statistical significance (p<4.3×10^−5^).

**Supplementary Figure 4**- Heatmap of partial genetic correlations for 16 traits including BP.

**Supplementary Table 1 –** Discovery and replication cohorts. **Supplementary Tables 2 –** Results of discovery and replication GWAS. **Supplementary Table 3 –** Results of COJO analysis for BP_ma_ GWAS.

**Supplementary Tables 4 –** Results of SMR-HEIDI for eQTLs and seventeen risk factors.

**Supplementary Table 5 –** Results of PhenoScanner v1.1 for three replicated loci.

**Supplementary Table 6 –** Results of genetic correlations for BP (LDHub).

**Supplementary Tables 7 –** Results of DEPICT analysis for SNPs with p<5×10^−8^ and p<1×10^−5^.

**Supplementary Tables 8 –** Results of MAGMA analysis.

**Supplementary Tables 9 –** Results of VEP and PAINTOR.

**Supplementary Tables 10 –** Tables with correlated DHS sites for *SPOCK2* and *CHST3* genes.

## ACKNOWLEDGEMENTS

This study was supported by the European Community’s Seventh Framework Programme funded project PainOmics (Grant agreement # 602736). The research has been conducted using the UK Biobank Resource (project # 18219). We are grateful to the UK Biobank participants for making such research possible.

The development of software implementing SMR/HEIDI test and database for GWAS results was supported by the Russian Ministry of Science and Education under the 5-100 Excellence Program”.

Dr. Suri’s time for this work was supported by VA Career Development Award # 1IK2RX001515 from the United States (U.S.) Department of Veterans Affairs Rehabilitation Research and Development Service. Dr. Suri is a Staff Physician at the VA Puget Sound Health Care System. The contents of this work do not represent the views of the U.S. Department of Veterans Affairs or the United States Government.

Dr. Tsepilov’s time for this work was supported in part by the Russian Ministry of Science and Education under the 5-100 Excellence Program.

We thank Eugene Pakhomov for developing software and database for eQTL-related analyses and Dr Sodbo Sharapov for assistance with data submission.

Cohorts for Heart and Aging Research in Genomic Epidemiology (CHARGE) Musculoskeletal

Working Group: We acknowledge the following individuals from the CHARGE Musculoskeletal Working Group as non-author contributors involved in the meta-analysis of data from CHARGE

cohorts : Cindy G. Boer, Michelle S. Yau, Daniel S. Evans, Andrea Gelemanovic, Traci M. Bartz, Maria Nethander, Liubov Arbeeva, Tuhina Neogi, Archie Campbell, Dan Mellstrom, Claes Ohlsson, Lynn M. Marshall, Eric Orwoll, Andre Uitterlinden, Jerome I. Rotter, Gordan Lauc, Bruce M. Psaty, Magnus K Karlsson, Nancy E. Lane, Gail Jarvik, Ozren Polasek, Marc Hochberg, Joanne M. Jordan, Joyce B. J. Van Meurs, Rebecca Jackson, Carrie M. Nielson, Braxton D. Mitchell, Blair H. Smith, Caroline Hayward, and Nicholas L. Smith.

## AUTHOR CONTRIBUTIONS

MF and YT contributed to the design of the study, carried out statistical analysis, produced the figures, and first draft of the manuscript; LC provided statistical and computational support; MP and PS analysed CHARGE dataset and contributed to interpretation of the results; YA and FW conceived and oversaw the study, contributed to the design and interpretation of the results; all co-authors contributed to the final manuscript revision.

## DATA AVAILABILITY

Summary statistics from our GWAS discovery and meta-analysis are available for interactive exploration at the GWAS archive (http://gwasarchive.org). The data set was also deposited at Zenodo (https://doi.org/10.5281/zenodo.1319332). The data generated in the secondary analyses of this study are included with this article in the supplementary tables.

